# IMAGENE: Single-cell association of live cell imaging and gene expression profiles of non-adherent cells through photoactivatable adhesives

**DOI:** 10.64898/2026.01.20.700510

**Authors:** Maria Isabella Gariboldi, Chloé Sturmach, Solène Bernard, Christian Weber, Maël Gourves, Yuki Umeda, Xueyang Li, Shinya Yamahira, Julien Fernandes, Asier Sáez-Cirión, Satoshi Yamaguchi, Florian Muller

## Abstract

Live cell imaging is uniquely placed to study cell behavior as it preserves spatial context and enables non-destructive observations over time. Integrating live cell imaging and molecular phenotypes with single-cell resolution is key to uncovering the relationship between the behavioral and morphological signatures of cells, and their molecular states. Non-adherent cells – as are most immune cells – however, present unique challenges in linking live cell imaging and fixed cell assays with single-cell resolution due to the difficulty of identifying individual cells across experimental modalities. To overcome this issue, we developed IMAGENE, an experimental and computational pipeline that leverages previously reported photoactivatable biocompatible adhesive material (PA-BAM) coatings for on-the-fly cell immobilization. We demonstrate the IMAGENE experimental and computational pipeline by generating a dataset of label-free time-lapse videos of primary human naïve CD8+ T cells following 24 hours of polyclonal stimulation. Individual cells, including highly motile cells, can be matched to expression profiles of genes of interest obtained through KrakenFISH, a modified version of the previously reported autoFISH setup for automated, single-molecule fluorescence *in situ* hybridization (smFISH) experiments that supports sample parallelization. We use this data to train explainable machine learning models that predict expression levels of individual genes, with variable performance, from hand-crafted dynamic and spatial features obtained from live cell imaging.

## INTRODUCTION

Immune cells display remarkable diversity in their morphology and behavior,^1,2,3^ features that are tightly coupled to their functional “state,” whether defined by phenotype or disease context.^4,5^ These physical and behavioral properties are controlled by molecular programs of cells and are critical determinants of the efficacy of the immune response or of therapeutic performance. For example, cell motility directly influences trafficking and tumour infiltration - key obstacles that currently limit the efficacy of CAR T-cell therapies in solid tumours.^6,7^ Despite their importance, these dynamic features remain difficult to characterise with conventional cytometric or *omics* approaches, which typically provide high-dimensional snapshots but lack temporal resolution. In contrast, live cell imaging uniquely enables direct, time-resolved measurement of dynamic cell morphology and behavior at single-cell resolution.^8,9^ Integrating molecular information with dynamic physical and behavioral characteristics of cells obtained through live cell microscopy supports the generation of models for non-invasive molecular profile prediction from live cell imaging as well as the study of the relationships underlying molecular and phenotypic traits.

Technologies linking live cell imaging and molecular profiling have gained increasing interest in recent years.^10,11,12^ A compelling example of such an approach is Raman2RNA, developed by Kobayashi-Kirschvink and colleagues^12^ to predict RNA profiles of cells from Raman microscopy data. As part of their work, they generate live-cell Raman measurements and fixed-cell RNA imaging data for a subset of cells, which are used to train high performing “anchored” models. However, their approach is not scalable to non-adherent or loosely adherent cells, as these cell types cannot be re-identified across experimental modalities, being washed off from substrates upon fixation and subsequent buffer exchanges. Approaches that adhere T cells - such as the use of polylysine or bioactive coatings (such as adhesion proteins) - significantly affect T cell mobility and phenotype, thereby limiting the ability to apply predictive models in common culture conditions. On the other hand, methods that physically confine individual cells, such as nanowell-based platforms, are compatible with non-adherent cells but inherently restrict cell migration and prevent interrogation of interactions between different cells.^13^ Developing methodologies compatible with non-adherent cells that overcome these limitations is crucial. Immune cells such as T cells offer an attractive model for such technologies because they play a major protective role in response to infection or vaccination, are central to contemporary cancer therapies, fundamental to immunological research, and display marked phenotypic and functional heterogeneity within and across donors.

To address this issue, we developed IMAGENE (Fig. 1), named based on its ability to correlate images to genes, an experimental and computational pipeline for integrating label-free time-lapse imaging data with gene expression profiles with single-cell resolution. The IMAGENE approach is enabled by a photochemistry system based on a photoactivatable biocompatible adhesive material (PA-BAM). This photo-responsive coating consists of a modified poly(ethylene glycol) (PEG) lipid bearing two fatty acid chains. Upon irradiation, the material loses one fatty acid chain and converts to a single-chain lipid, which results in cell-type agnostic cell attachment mediated by the lipid’s interaction with cell membranes. Previous work showed the use of this coating for versatile multi-cell type patterning applications by alternating cycles of structured light irradiation and cell suspension incubation.^14,15^ In this work, we show that this coating also permits on-the-fly immobilization of cells and that the resulting fixed cells can be reidentified and characterized through quantitative fixed cell assays such as single molecule fluorescence *in situ* hybridization (smFISH). A graphical visualization of our method can be seen in <u>Supplementary Video 1</u>.

**Figure 1.**
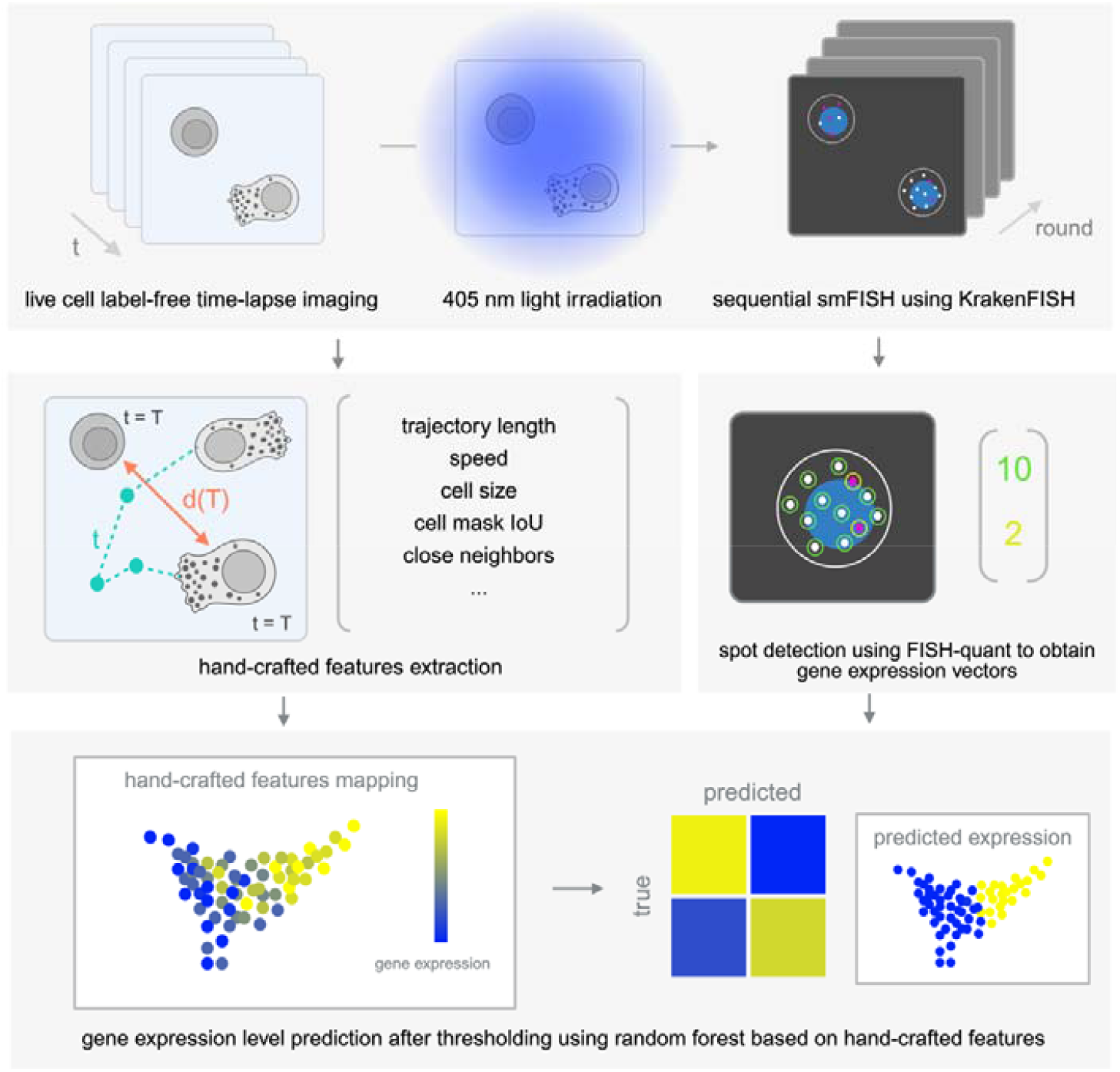
IMAGENE overview. Cells are imaged non-invasively through live time-lapse imaging, immobilised *in situ* on-the-fly using 405 nm light irradiation and their gene expression patterns are probed through smFISH. Hand crafted features are extracted from label-free images to quantify dynamic morphology, trajectory patterns and neighbourhood information. RNA expression for genes of interest is quantified using FISH-quant. Hand crafted features quantified from live cell images and RNA expression levels can be mapped across data modalities and explainable models are developed for the prediction of gene expression profiles from live cell imaging features.

To support programmed processing of several samples in parallel, we developed KrakenFISH, a modified version of our previously reported autoFISH system that supports automated, sequential smFISH experiments.^16^ IMAGENE enables low-cost experimental iterations and is compatible with a broad range of imaging setups and downstream fixed cell characterization approaches. Furthermore, its design permits training models that can be deployed in common culture conditions. We demonstrate the use of IMAGENE by generating a dataset of label-free time-lapse images of live human CD8+ T cells after 24 hours of polyclonal stimulation matched at single-cell resolution with “molecular ground truth” labels consisting of expression profiles of a set of genes of interest. We chose a well-established biological model to support careful monitoring of cell reidentification performance across modalities. While the starting pool of naïve CD8+ T cells is homogenous, the gene expression, the physical appearance and the behavior of cells within 24 hour stimulated samples are remarkably diverse. We show the approach’s ability to track features related to cell morphology, trajectory and neighbourhood from label-free time-lapse images and use the resulting dataset to train machine learning models to predict the expression level of individual genes from these live cell features.

## RESULTS

### Photoactivatable adhesives allow on-the-fly immobilization of motile cells

The experimental pipeline underlying the IMAGENE platform critically relies on the ability to dynamically tune cell-substrate interaction of non-adherent cells to enable, on one hand, non-invasive imaging of unbound live cells, and on the other, RNA imaging of adhered fixed cells. We achieve this transition by leveraging an advanced material coating to induce *in situ* “capturing” of cells following live cell imaging. Specifically, this photoactivatable biocompatible adhesive material (PA-BAM) can be triggered using 405 nm or near-UV light. In its native state, the hydrophobic chains of the PA-BAM coating self-assemble into nanoscale structures that were previously shown to cooperatively inhibit cell anchoring.^14^ Upon irradiation with 405 nm light, the modified PEG undergoes dissolution of its self-assembled structure, resulting in cell attachment through lipid-cell membrane interactions (Fig. 2a). Following irradiation, the material converts from a dual chain lipid into a single chain lipid structure by loss of a photocleavable chain (Fig. 2b). Previously reported uses of this material consisted of iterations of micro-patterned activation of the substrate followed by cell suspension incubation to achieve fine single-cell patterns. The material was shown in these studies to be highly biocompatible, as cells retained viability even after hours of incubation.^14^

**Figure 2.**
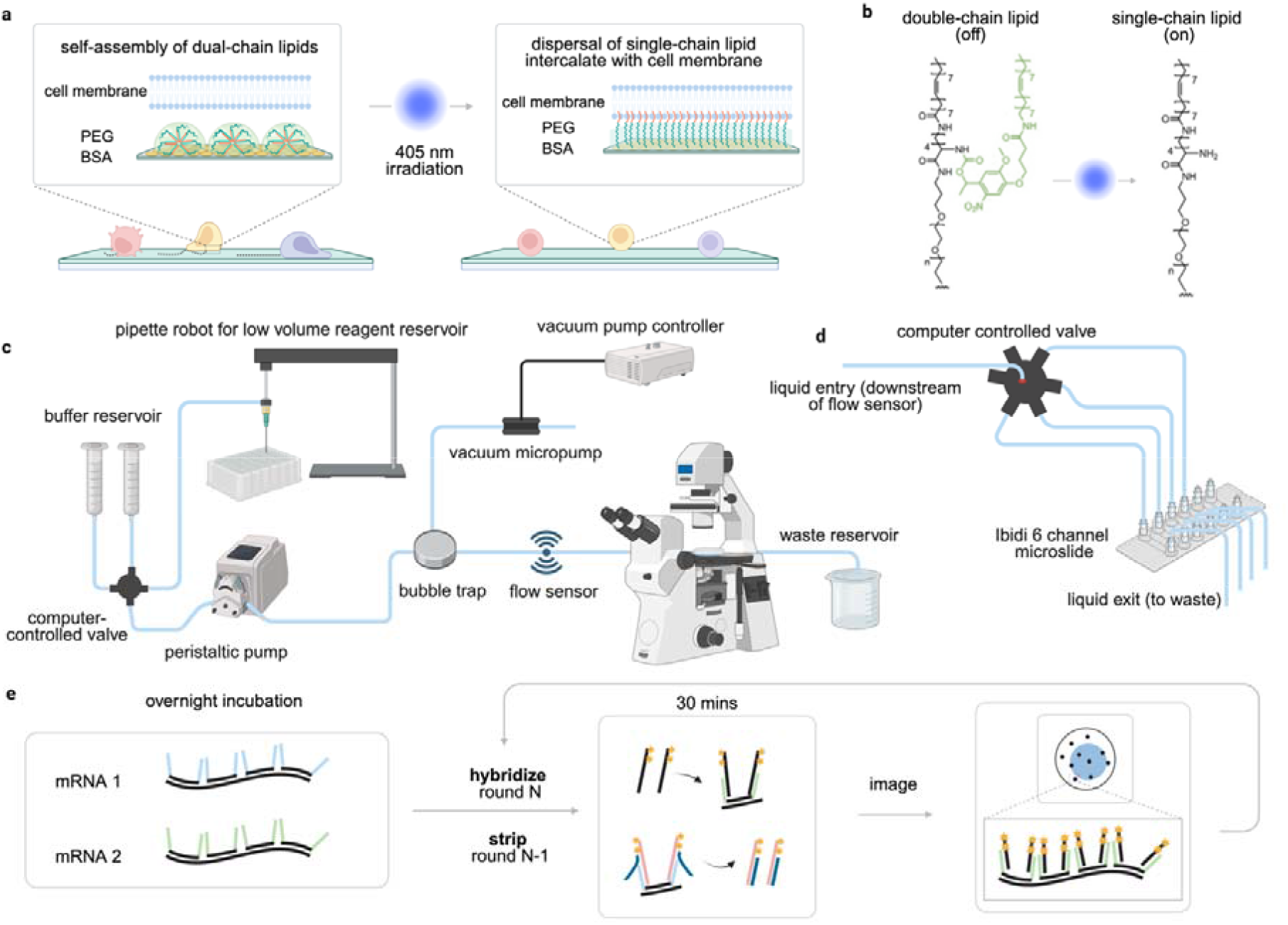
IMAGENE experimental pipeline for sequential live cell and smFISH imaging of the same cells. (a) Label free live cell imaging is performed with PA-BAM coating in its native state, which consists in dual chain lipids self-assembled into nano-scale structures that cooperatively inhibit cell attachment. Upon irradiation, loss of an oleyl group results in conversion from a two-tailed PEG to a single-tailed PEG that can intercalate cell membranes, stably anchoring cells to the substrate. (b) Photochemical reaction resulting in loss of an oleyl group and conversion of dual chain lipid into single chain lipid structure. (c) Previously reported AutoFISH open-source, flexible fluidic setup for low-cost smFISH experiments. (d) Modification of fluidic system through an addition of a computer-controlled valve and consumable microchannel slide support (Ibidi) downstream of flow sensor to obtain KrakenFISH, a modified version of AutoFISH for parallel assessment of up to six experimental conditions. (e) autoFISH and KrakenFISH simplified modular smFISH chemistry approach representation. Primary probes complementary to genes of interest are incubated overnight and contain barcode “flap” sequences specific to each RNA species. Each gene can then be iteratively hybridized and imaged thanks to secondary fluorophore-conjugated probes complementary to barcode regions. For each round, secondary probes from the previous round can be simultaneously stripped during hybridization through the use of displacement probes. Panels a and b are adapted from Yamahira et al. (2022)^14^ and panels c and e are adapted from Weber et al. (2025).^16^

We hypothesised that the same material could be used to achieve on-the-fly immobilization of cells, including highly motile cells, incubated directly on the coating. We therefore used an empirical approach to optimize illumination parameters – namely illumination intensity and exposure time - to be able to capture cells in place directly on the microscope (Supplementary Fig 1). At optimal parameters, we achieved good efficiency of cell anchoring. We found that parameters for cell capture should be optimized for each imaging system and updated regularly. For our system, we found that illumination of 2 seconds at 50% intensity with 405nm wavelength on an LED8 (Leica microsystems, Germany) light source for each field of view were sufficient to obtain reproducible attachment of cells using a 40x, 1.3 NA oil immersion objective, while ensuring attachment was localized to the field of view with no significant unspecific cell attachment.

### KrakenFISH automates multi-condition sequential smFISH

Once we demonstrated the ability to perform light-triggered immobilization of cells directly on the microscope, we built an experimental pipeline for generating datasets coupling live cell time-lapse imaging data with gene expression profiling obtained through sequential smFISH for the same set of cells. Sequential smFISH is a highly quantitative technique in which rounds of probe hybridization and imaging are alternated to visualize multiple RNA species, with each RNA molecule appearing as a diffraction-limited spot

To automate sequential smFISH experiments for several samples in parallel, we developed KrakenFISH, a modified version of our previously reported autoFISH system (Fig. 2c), an open-source fluidic set up that can be mounted onto fluorescence microscopes for low-cost, customizable, sequential smFISH.^16^ The modification of the original autoFISH setup consisted in adapting the fluidic system to be compatible with multi-channel microfluidic microslides using Luer connectors and the addition of a computer-controlled valve to precisely control fluid exchange in downstream microchannels (Fig. 2d). This modification enabled assessing several conditions in parallel, increasing the throughput of experimental conditions up to 6-fold (based on the number of channels in standard Ibidi microchannel slides). The system uses the same modular smFISH chemistry approach previously described by Weber et al. (2024).^16^ (Fig. 2e), which was optimised to maintain low recurrent experimental costs. While we chose RNA imaging to generate quantitative molecular ground truth labels, we expect the IMAGENE approach is compatible with a variety of fixed cell quantitative imaging approaches, such as protein expression quantification through immunofluorescence (Supplementary Fig 2).

### IMAGENE reidentifies non-adherent cells across experimental modalities

To find corresponding cell pairs between live cell imaging data and smFISH images of anchored cells, we designed a computational pipeline described in Fig 3. Briefly, live cell image segmentations on final timepoints were obtained using Cellpose^17^ with manual quality control, followed by SAM2,^18^ to respectively obtain initial seeds and to back-propagate segmentations across all timepoints (Fig. 4a). Each located cell could then be re-segmented using a fine-tuned Cellpose model followed by feature extraction, with selected feature examples shown in Fig 4b-d. Segmentations for smFISH were also obtained semi-automatically using a finetuned Cellpose segmentation model with manual quality control. Spot detection was performed using FISH-quant^19^ to assign gene count vectors to individual cell segmentations (Fig 4e). Cell-reidentification between live and fixed-cell imaging was performed by identifying the optimal translational shift between the smFISH segmentation and the segmentation of the last timepoint of the live cell imaging that yielded the lowest cumulative distance between nearest cell pairs. This combinatorial enumeration approach yielded a clear minimum (Fig 4g), ensuring confident alignment between the two datasets.

**Figure 3.**
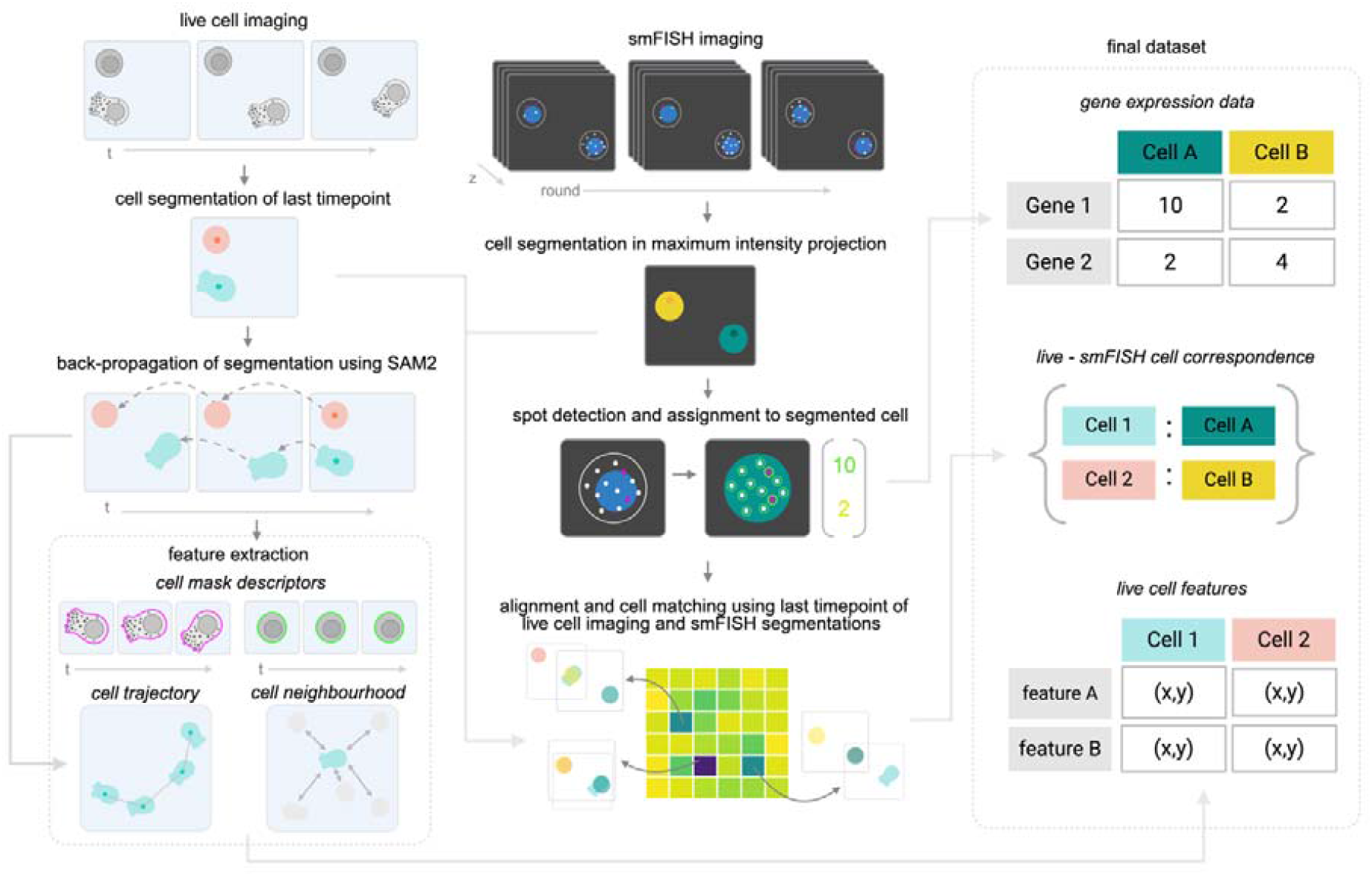
IMAGENE computational pipeline for matched dataset generation. The last time point of live cell images was segmented using Cellpose. Centroids were used as initial landmarks for SAM2 enabling to back-propagate cell segmentations across timepoints for single cells. Tracked cells were then re-segmented using a finetuned Cellpose model to obtain more precise contours. Trajectories and contour information was used to obtain features relating to morphology, trajectory and cell neighbourhood. For smFISH image processing, a maximum intensity projection of smFISH images was segmented using Cellpose and RNA spot detection was performed using FISH-quant followed by spot assignment to individual cell masks to generate gene expression profiles for each cell. Cells overlapping with bright artefacts were excluded from the analysis (not shown). The last frame of the live cell imaging time-lapse segmentation, obtained through SAM2, and the smFISH segmentation were aligned after magnification adjustment using a brute-force approach for finding the optimal relative translational shift between the two segmentations. This approach consisted in applying a sliding window to find the relative positioning yielding the minimum sum of distances of each segmented cell in the last frame of the live cell time-lapse to the nearest cell in the smFISH segmentation. Once the optimal translation was applied, cells were paired across datasets based on spatial proximity, generating a correspondence between unique cell identifiers of the live cell segmentation and the fixed cell segmentation. Cell pair filtering through d_1_ and d_2_ metrics is not shown.

**Figure 4.**
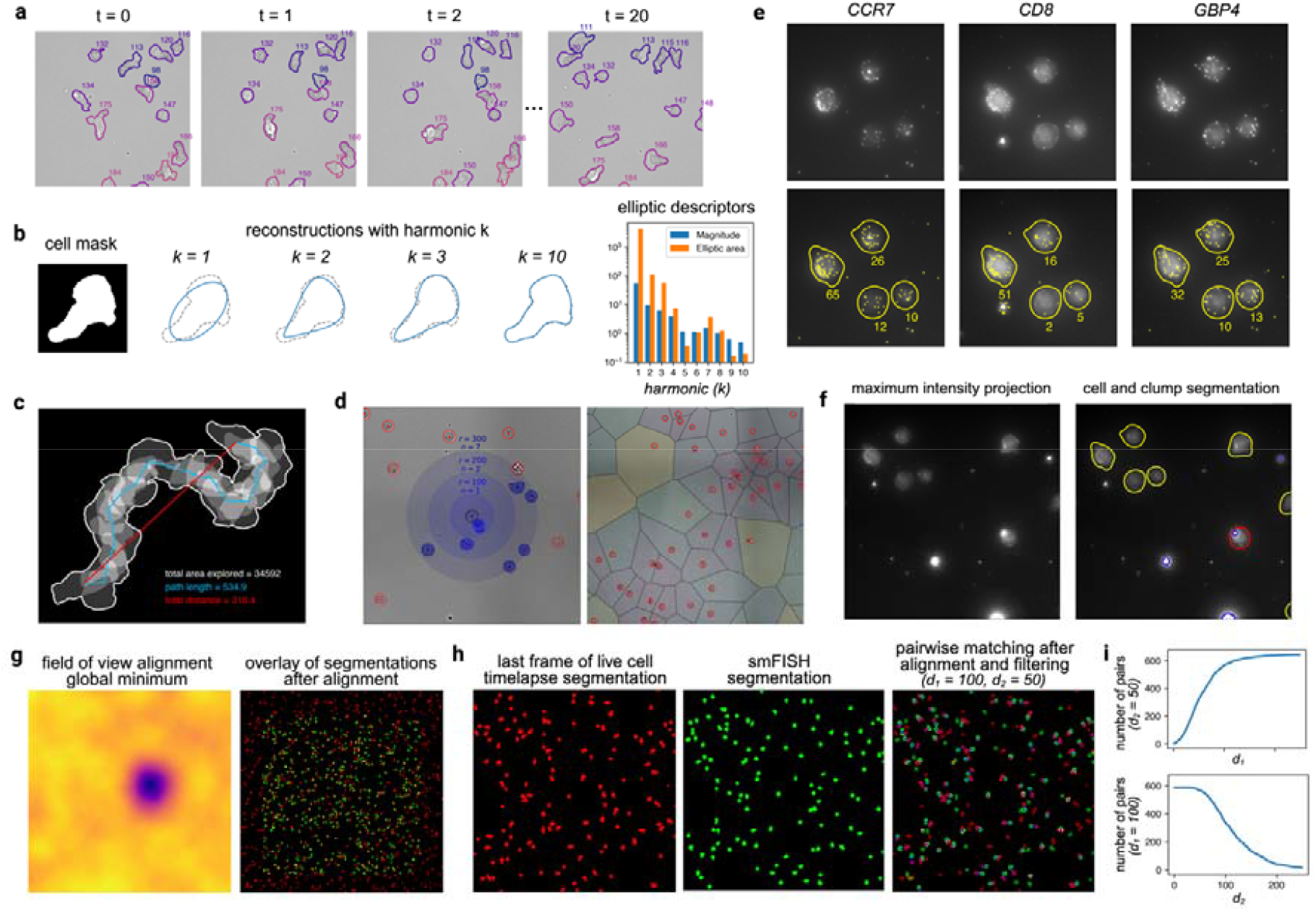
IMAGENE dataset generation results. Live cell time-lapse image processing approach and feature extraction (a-d). (a) Example result from backpropagation of cell segmentations using SAM2. Re-applying our finetuned model to propagated segmentations supports extraction of features related to cell contour at every timepoint, such as features computed from elliptical Fourier descriptors (EFDs). (b) Demonstration of EFD application to sample cell contour at different harmonics (*k*) and resulting magnitude and elliptical area measures. (c) Example features extracted from cell trajectory analysis such as total area explored, path length and total distance travelled. (d) Example of neighbourhood features, including neighbours within a given radial distance and Voronoi tessellation features. Fluorescent smFISH image processing approach and spot detection (e-f). (e) smFISH images are processed through (e) segmentation, spot detection and assignment to individual cell masks are shown for sample genes. (f) Bright artefact detection overlayed with cell segmentation and removed cells overlapping with bright unspecific signal. Detected cell outlines are shown in yellow, detected artefact outlines in blue, and cells removed due to artefacts are displayed in red. (g-i) Cross-modality single cell pairing results. Segmentations for the last frame of the live cell imaging and the smFISH are aligned through a brute-force approach enabling the identification of a global minimum distance across cell pairs. (g) Global minimum identification and corresponding cell segmentation alignment. (h) Cell pair matches can be dynamically filtered based on the distance to matched cell (d_1_) and distance to the second closest match cells (d_2_), measured in pixels. (i) Influence of decreasing d_1_ and increasing d_2_ in total number of cells in the final dataset.

With this pipeline, we were able to generate features for live cells, gene expression vectors for fixed cells as well as successfully reidentify a high portion of cells across experimental modalities. Recognising the trade-off between cell matching certainty and size of the matched dataset, we established two user-adjustable quality metrics (Fig 4h and i): the maximum distance between paired cells (d_1_) and the minimal distance to the second closest cell in the alternate modality (d_2_). This last metric was applied to the second closest smFISH segmented cell to a given brightfield segmented cell and vice versa. These metrics can help dynamically adjust confidence thresholds for matching based on visual inspection and variable experimental parameters such as cell density (Supplementary Fig 3).

Fig 5a shows the loss of cells at each step of the IMAGENE pipeline for one of our samples. Considering only the area of imaged live cells that overlapped with the area imaged for smFISH, 487 cells were present in the fixed cell modality as opposed to 528 cells in live cell modality. Without accounting for the possibility of new cells attaching to the material during washing, this suggests an estimated effective cell capture rate through the on-the-fly photochemistry system of >90%. We were able to generate matches with reasonable confidence in this area after applying d_1_ and d_2_ thresholding for 298 cells, that is 61% of total immobilised cells and 56% of total live cells in the imaged area. Through this filtering process we adopted a rigorous approach that favours confidence in the matches, accepting a smaller dataset as a consequence. While these results will vary depending on experimental parameters such as cell motility and density, we estimate that a good rule of thumb is to image in both modalities an area including at least twice as many cells as the desired dataset size.

**Figure 5.**
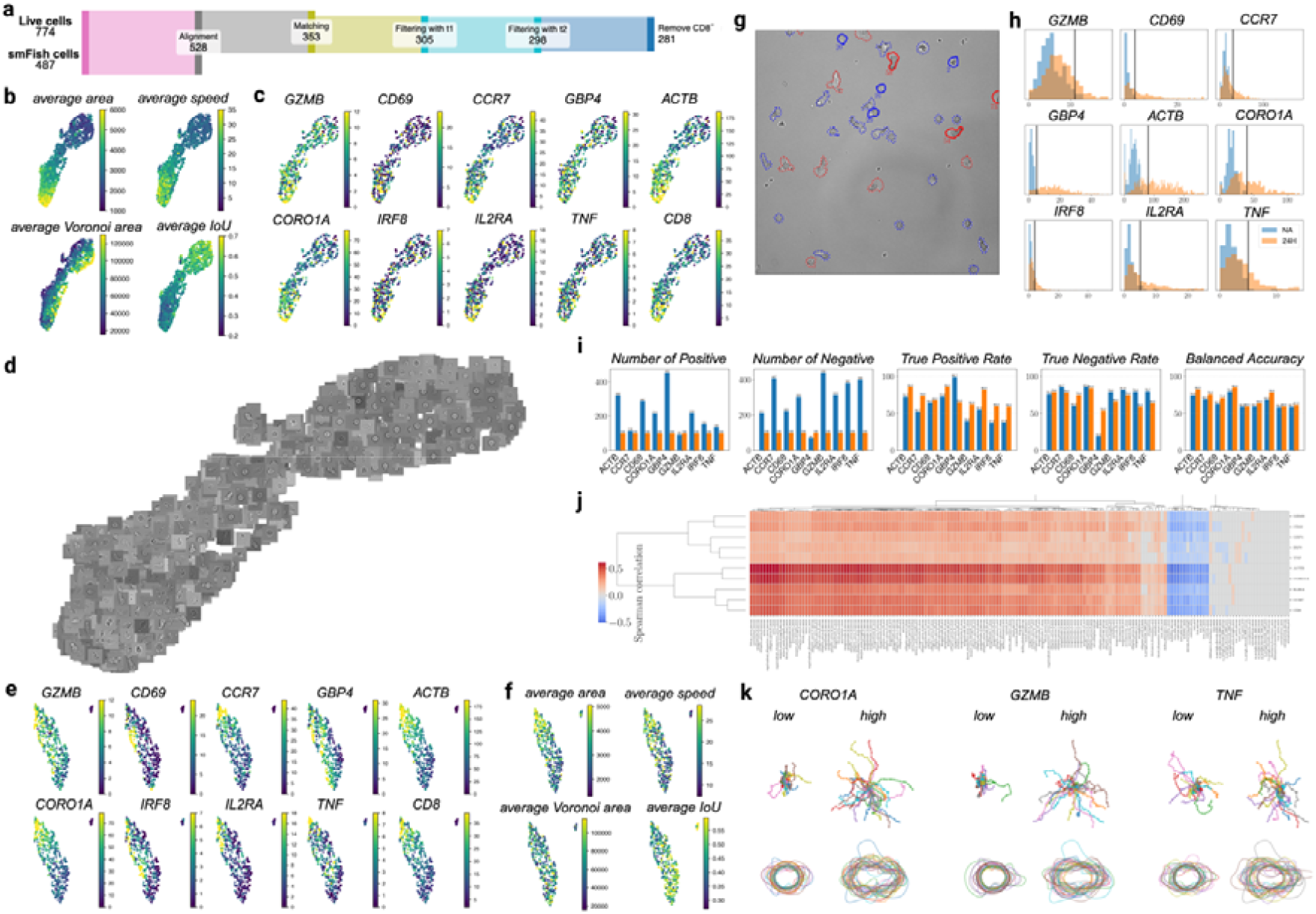
Integration of image-based feature extraction and gene expression profiling for stimulated naïve CD8 T cells. (a) Depiction of cell loss at main steps in the pipeline to generate the final cross-modality dataset for an individual sample. Note that only a subset of the area imaged in live modality was imaged for smFISH, this is captured by the alignment step, which only includes live cells present in the smFISH imaged area. (b) UMAP dimensionality reduction projection of live cell features colour-coded by different example features, and (c) the same projection applied to matched cells colour-coded by gene expression level. (d) Live cell feature UMAP projection overlayed with single timepoint image of each cell. (e) UMAP dimensionality reduction projection of genes assessed through smFISH color-coded by each gene assessed and (f) the same projection overlayed with selected live cell features for matched cells. (g) Example of deployment of model showing gene prediction as high (red) or low (blue) for CORO1A. Solid lines indicate cells for which the ground truth was available, with the actual value shown. Dotted lines are used for cells for which a high-quality match was not established. Note that for this visualization, training and test sets were not distinguished. (h) Threshold choice for gene expression profiles used to define gene expression as high or low in the threshold-based binary classification model in 24 hour activated cells (orange) relative to naïve baseline expression (blue). (i) Random forest model classification results for whole distribution thresholding of genes (blue) versus edge-case training (orange). (j) Live cell feature-gene expression correlation matrix and (k) overlay representation of cell contours reconstructed using elliptical Fourier descriptors (EFDs) and trajectories for top 30 and bottom 30 expressing cells for selected genes.

### IMAGENE links time-lapse image-derived fingerprints with gene expression profiles for single cells

To account for limitations in cell purity inherent to isolation kits, cells not expressing at least two transcripts of the CD8 marker gene were excluded from the analysis. To obtain a more precise contour estimation of each cell for morphological feature extraction, we trained a Cellpose model on manually annotated data to precisely track cell contours, including cell protrusions and fine lamellipodia structures. After application of SAM2, this model was reapplied to cells across timepoints to combine SAM2’s ability to track cells, providing spatial trajectory information, with our finetuned model’s precision in contour detection (Fig. 4a). From the combination of cell contour data and their location in space at any given timepoint, we extracted a total of 163 features that describe cell outline (Fig 4b), migratory behavior (Fig 4c) and neighbourhood (Fig 4d). A complete list of extracted features can be found in Supplementary Table 1.

Cell segmentation and RNA detection using FISH-quant^19^ were combined to assign transcript counts to each cell for each gene (Fig 4e and g). As a quality control step in the smFISH data analysis, we identified aggregates and autofluorescent debris through manual thresholding of smFISH images. Cells overlapping with these aggregates were excluded from the analysis (Fig 4f).

This pipeline allowed us to generate datasets containing on one hand rich information of live cell behavior, and on the other hand gene expression vectors for each cell, with a high-fidelity subset of cells from each dataset being identifiable across modalities. To demonstrate the applicability of models trained on data generated using the IMAGENE pipeline to common culture conditions and confirm the specific PA-BAM substrate does not affect cell morphology and behavior, we compared the distribution of features extracted from cells imaged using IMAGENE to cells imaged on other substrates. Unlike polystyrene cell culture labware, we found that CD8 T cells non-specifically attach to uncoated glass over time (data not shown). We found that low adhesion substrates such as Ibidi Bioinert microchannel slides resulted in comparable cell feature mapping (Supplementary Fig. 4), suggesting a model trained using IMAGENE should be deployable to make predictions of cells imaged on any low attachment, inert substrate.

### Predictive performance of models trained on CD8 T cell dataset is gene-dependent

To demonstrate the pipeline’s ability to identify associations between live cell features and gene expression profiles within individual samples, we applied IMAGENE to naïve CD8 T cells after 24 hours of stimulation. We found that combining several treatment conditions (e.g. stimulation levels) could result in potential over-estimation of the pipeline’s performance, as even random matching within each sample could enable observing correlations between morphology and gene expression levels if both morphological features and gene expression distributions are affected by the treatment condition. We therefore chose to focus on intra-sample variation and selected a well-accepted model of morphological transition to ensure that our technology could be evaluated against easily interpretable, benchmarked cellular responses. We therefore selected polarization of naïve CD8+ T cells as a model, as their morphological transitions, particularly their increase in size, following polyclonal stimulation are a well-established phenomenon.^20^ We applied the IMAGENE pipeline to cells following 24 hours of stimulation and assessed their gene expression changes relative to cells maintained in culture without stimulation.

Starting with 1,545 live cells and 1,008 cells assessed through smFISH across two samples (sample replicates both contained cells after 24 hours of polyclonal stimulation), we were able to obtain 540 matched cells (respectively 281 and 259 in each sample) after excluding cells not expressing our CD8 cell type marker, as previously described. Ten genes were included in the analysis out of the 12 probed using smFISH (IFNG and XP01 were excluded due to poor smFISH quality in this experiment). CD8 was included as a cell type marker, CCR7 due to its recognition as a naïve/memory T cell marker; ACTB and CORO1A because of their involvement in cytoskeletal remodelling; CD69, GZMB, IL2RA, and TNF because of their roles in activation and effector functions; and GBP4 and IRF8 based on their use in optimization experiments displaying distinctive expression patterns (Fig. 5e). Along with the two samples used for the analysis, a sample of non-stimulated cells was immobilised and probed using smFISH to help establish baseline expression levels of genes. Despite starting from a homogenous population of naïve cells, remarkable diversity arises in both the physical features of cells and their gene expression profiles (Fig 5b-f). Unsurprisingly, many of the genes probed showed a high degree of correlation, being largely associated with activation and actin remodelling.

The IMAGENE pipeline enables generating population maps to visualise the correlation of different live cell features with each other and with genes of interest (Fig 5b-f). We next explored the ability of IMAGENE to develop predictive models for gene expression levels. To ensure model interpretability, we trained two types of random forest models taking as input live cell features to predict gene expression levels as high or low for each gene. The first type labelled cells as high or low based on thresholds set manually while contextualising expression levels to non-stimulated cells (Fig 5h). The second model was trained using respectively the top and bottom 100 expressing cells. Edge-case training led to improved classification for all genes demonstrating the first model was likely affected by many cells having expression levels close to threshold values (Fig 5i). While ACTB, CCR7, CORO1A, IL2RA achieved relatively high performance (78.5-85% balanced accuracy in edge case training), CD69, GBP4, GZMB, IRF8 and TNF performed higher than random but still below 70% balanced accuracy, even when adopting an edge-case training approach. We can visualise feature associations with genes of interest using a correlation matrix (Fig 5j) and by overlaying trajectory and boundaries for cells at the edges of the distribution (Fig 5k). This visualization approach enables concluding, for example, that for some genes such as CORO1A and GZMB, edge cases have markedly different cell trajectories, whereas the same pattern is not as striking in high versus low TNF-expressing cells. To visualize how a similar model could be deployed on cells in culture, we applied the model to all live cells in a selected time-lapse area, including cells for which a molecular ground truth was not available (Fig 5g).

The differential performance of trained models depending on genes suggests that low-performing genes don’t produce clearly discernible phenotypes as assessed through imaging, at least in the biological conditions and at the levels assessed, or that the simple set of features employed are not sufficiently discriminating for these genes. Class imbalance was also gene-dependent, and the lack of representation of one of the classes might explain the low predictability of some genes. For example, only 68 cells in the whole dataset were classified as having low expression of GBP4, resulting in a particularly small representation of this gene in the test set. Features related to cell morphology and trajectory were more predictive across different genes compared to those describing cell neighbourhood.

## DISCUSSION

We developed the IMAGENE experimental and computational pipeline to map non-adherent cells across live cell and fixed cell imaging modalities. The IMAGENE pipeline leverages a previously reported biocompatible photochemistry system that we show can also be applied to immobilize cells *in situ* on-the-fly. We show that through a “brute-force” approach, we can find an optimal alignment of cells by minimising the collective distance across cell pairs, and we can further enforce quality matching through user-defined metrics. We demonstrate this approach by applying IMAGENE to identify features of human naïve CD8+ T cells after 24 hours of polyclonal stimulation associated with gene expression patterns linked to cell activation. Pooling matched cells from different experimental conditions could be misleading as it may generate interpretable results even if random matching occurred within each sample, if both gene expression and cell morphology distributions are shifted in treatment conditions. For this reason, we focused on identifying signatures associated with intra-sample variation to provide confidence that IMAGENE correctly reidentifies individual cells across imaging modalities. These findings, in turn, demonstrate the utility of this single-cell approach, as rich information can be extracted by exploiting natural variation of genes within individual samples, not just across different perturbation conditions.

A few tools have been developed in recent years to support mapping between live features of cells and their molecular profiles. For example, a notable approach that addresses a similar question as IMAGENE is the robotic integration of imaging and single-cell RNA sequencing using cell pickers reported by Jin and colleagues.^21^ This approach has considerable advantages, especially in terms of depth of sequencing and not requiring *a priori* gene candidates, relying on next generation sequencing approaches. On the other hand, the solution may be less flexible and accessible, given the significant hardware component, and less scalable in terms of number of cells (authors cite cell isolation times of roughly 15 minutes for 96 cells). Unlike flow-based and spatial confinement methods, IMAGENE can be applied to cells incubated on any glass bottom substrate, thereby supporting the production of data to train models that are easily deployable in common-culture conditions as well as image-based high throughput screening applications. Furthermore, IMAGENE has no specific hardware requirements. If small sets of genes or proteins are probed, cells can be analysed without our automated KrakenFISH system, rendering IMAGENE easily accessible without the need for specialised equipment other than a fluorescent microscope.

Another significant advantage of IMAGENE is its flexibility. For example, it is compatible with other quantitative imaging approaches, including immunofluorescence. IMAGENE is also integrable with other live cell imaging approaches - both label-free and fluorescent – provided that illumination does not extend into the near-UV range. On the other hand, choice of fluorescent channels must take into consideration downstream fixed cell analysis methods, although chemical or photo-bleaching approaches prior to, for example, smFISH may help overcome this constraint. Further, the photochemistry of the cell capture system relies on lipid bilayer interactions, rendering it compatible with virtually any suspension cell type. Another significant advantage is the ability to decouple live cell and fixed cell experimental modalities, i.e. to conduct live cell imaging and fixed cell imaging on different systems and at different times, while bearing in mind considerations related to RNA degradation. This provides substantial flexibility when working, for example, with infectious samples. In this case, live cell imaging can be performed on any imaging system available in a high-biosafety environment, provided a 405-nm excitation source is available. Subsequent processing can then be carried out after fixation on appropriate equipment. Finally, acquiring phenotypic and molecular data in two separate steps also renders the technique compatible with assessing live cell features associated with intracellular markers, notably transcriptional hubs which play a key role in driving CD8+ T cell fate.^22^ In fact, markers that are not localized on the cell membrane require cell fixation and permeabilization to be probed, thereby not being associable to live cell modalities when using “one-shot” technologies such as imaging flow cytometry and ghost cytometry^23^ approaches.

It is important to keep in mind some crucial limitations in interpreting data from IMAGENE. Every step of the pipeline comes with a certain level of uncertainty. Accordingly, the model’s performance captures both the underlying informativeness of the cellular features and the reliability of labels produced through a multi-stage process. For example, smFISH spot detection is inherently susceptible to error, particularly with very low expressing genes, whose quantification is more vulnerable to small artefacts and noise, and genes that come close to spatial saturation, which may result in the inability to discern individual spots. Cell pair matching is also imperfect, particularly as many cells can still move some distance before becoming completely immobilised. Matching is particularly challenging in high density areas. Our matching stringency metrics may thus need to be revisited in situations where cell clustering is an interesting feature. In our case, we did not find a relationship between the genes we probed and features describing cell neighbourhood. Nonetheless, these features may be of great importance for other biological models including co-culture systems to study interactions between different cell types, as well as many mono-culture systems, since paracrine signalling, quorum sensing and swarm behavior have been demonstrated to be relevant phenomena in many immune cell models.^24,25^

While we focused on intra-sample variation to provide high confidence in our cell re-identification approach, IMAGENE can be scaled to higher throughput formats within a set of essential constraints. For example, the IMAGENE pipeline can be carried out in open multi-well formats, enabling the molecular ground truth generation to be compatible with high content screening and robotic solutions for fluid handling for smFISH or other molecular profiling approaches. However, while a benefit of our approach is the ability to image several cells simultaneously through tiled time-lapse acquisitions, a crucial parameter that may limit the tiled acquisition size is the time between imaging and light-induced capture of cells in the last field of view, to prevent large cell displacements prior to immobilization affecting matching performance. This will require optimization of acquisition-capture-fixation routines. Furthermore, back-to-back rounds of imaging and immobilization of cells are supported by experiments conducted on BaF3 cells demonstrating high viability even after days of being immobilised on the coating (Yamahira et al., unpublished manuscript). However, if cells are immobilised for prolonged periods as part of the IMAGENE pipeline, appropriate smFISH controls would be advisable to ensure genes of interest are not affected. Another important parameter to control for is cell density. In optimising our system, we found that our combinatorial approach to matching failed to identify a clear minimum if either the efficiency of cell immobilization was too low or the total cell density was too high.

When applied to stimulated naïve CD8+ T cell samples, our approach could not clearly establish specific morphological signatures uniquely associated with individual genes, given the high correlation between selected genes under polyclonal stimulation conditions. For this proof-of-concept, we focused on a subset of well-established genes. For follow up studies, larger gene panels could be employed. These could be integrated by either increasing the number of imaging rounds, or implementing multiplexed detection schemes as used in spatial transcriptomics. We chose to implement a compact model, to ensure interpretability of results, on a well-established biological system, to guarantee cell-reidentification performance could be evaluated with confidence. A logical extension of our approach is through the integration of more nuanced phenotypes by addition of texture features, such as features related to granularity, as well as through the integration of deep learning approaches, such as convolutional neural networks or transformer-based approaches. While we expect texture features could have improved model performance, we found these metrics to be highly sensitive to batch effects as well as local variations in focus and illumination. Nonetheless, use of these metrics is reported in the literature,^26^ suggesting that integration of such features, as well as deep learning approaches, is highly plausible, provided that these variations are systematically accounted for experimentally and computationally.

In conclusion, we show IMAGENE enables linking live cell and molecular features of non-adherent cells through photoactivatable adhesives. This molecular ground truth approach can be used to both explore the interplay between molecular and biophysical properties of cells, as well as to deploy predictive gene expression profiling models in common culture conditions. Being a single cell approach, IMAGENE is well placed to not only assess the influence of chemical and genetic perturbations in different samples, but also to explore relationships between phenotype and molecular profiles leveraging natural variation in gene expression. Furthermore, this pipeline could be used in combination with a smart microscopy system and a cell picker or digital micromirror device to enrich for specific cell populations. For example, live cells predicted to express genes of interest based on signatures extracted from images could be immobilised or isolated. This approach could be of particular interest for generating predictive models for intracellular markers, which are difficult to employ for isolating live cells. Altogether, we envision IMAGENE could be impactful in a variety of applications based on profiling of non-adherent cells - particularly immune cells - including drug development, digital integration in biomanufacturing, diagnostics and personalised medicine.

## METHODS

### Photoactivatable coating preparation

Photoactivatable biocompatible adhesive material (PA-BAM) was synthesised as previously reported.^14^ Phosphate-buffered saline (PBS) without calcium or magnesium was used for all sample preparation steps. Ibidi glass-bottom slides containing six microchannels were washed with PBS and coated overnight at room temperature with a 1% solution of fatty acid-free bovine serum albumin (BSA) (Sigma-Aldrich). Channels were then thoroughly washed with PBS to remove any excess BSA. Channels were incubated with 125 μM PA-BAM in 50% DMSO in PBS and incubated at 37°C in a humidified atmosphere for three hours. Finally, channels were washed thoroughly with PBS and stored at 4°C protected by light.

### Isolation and *in vitro* stimulation of primary human CD8+ T cells

Samples from blood donors were obtained from the Établissement Français du Sang in the context of a collaboration agreement with the Institut Pasteur. Peripheral blood mononuclear cells (PBMCs) were separated from peripheral blood by density gradient centrifugation. Naïve CD8 T cells were then isolated from buffy coats using EasySep Human Naïve CD8+ T Cell isolation kit (STEMCELL Technologies) according to manufacturer’s instructions. Cells were grown in phenol red-free, serum-free medium ImmunoCult (STEMCELL Technologies) supplemented with interleukin-2 (Sigma-Aldrich) at 110 IU/mL. The choice of medium was dictated by the interference of serum with immobilization of cells using PA-BAM coating. Cells were stimulated for 24 hours or kept in non-activated conditions. Stimulation was performed by placing cells at 1 million cells/mL in wells pre-coated overnight with anti-human CD3 (Invitrogen) and anti-human CD28 (Invitrogen) antibodies, respectively clones OKT3 and CD28.2, both diluted to a final concentration of 10 μg/mL in PBS.

### Live Cell Label-free Imaging and photoimmobilization

Following stimulation, cells were resuspended and placed in glass bottom six-channel microchannel slides (Ibidi) coated with PA-BAM. Live cell microscopy imaging was performed on a Thunder Imager (Leica microsystems, Germany) epifluorescent microscope using a 40x NA 1.3 oil immersion objective and LED8 (Leica microsystems, Germany) illumination source. Live cell imaging conditions were maintained using an Okolab live cell imaging chamber (Okolab, Italy). Cells were left to equilibrate on the microscope for at least 30 minutes. Tiled time-lapse image acquisitions were obtained in two channels for 24 hour activated cells, and in one channel for non-activated cells to provide baseline gene expression levels. Cells were imaged as a 4x4 tiled acquisition across four focus planes (1,5μm between consecutive planes) for 21 time-frames, with an optimised time interval of 21 seconds between consecutive time-frames, for a total time-lapse length of 7 minutes, 21 seconds. At the end of the live cell time-lapse acquisition, each tile was illuminated with 405 nm light at 50% LED power for 2 seconds using the LED8 (Leica microsystems, Germany). After the acquisition and immobilisation was completed for each condition, cells were incubated for at least one minute prior to moving to the next condition to ensure stable cell anchoring. After all conditions were acquired, cells were washed with PBS, fixed in a 4% paraformaldehyde solution for 30 minutes and washed thoroughly with PBS.

### Fixed cell preparation for KrakenFISH

After fixation, cells were washed twice with 2X SSC (in water containing diethyl pyrocarbonate, DEPC) and permeabilized with 0.1% Triton-X-100 in 2X SSC for 10 minutes at room temperature. Channels were washed twice with 2X SSC for five minutes, and then washed twice with 20% formamide in 2X SSC for 8 minutes. Finally, cells were incubated overnight in a humidified chamber at 37 °C in the primary hybridization buffer, consisting of primary oligos at a final concentration of 100nM each, in 2X SSC, 20% Formamide, 10% Dextran. After primary hybridization, channels were washed four times for 15 minutes at 37 °C with 20% formamide in 2X SSC, followed by a wash in 2X SSC and then mounted on KrakenFISH for subsequent washes and secondary hybridization steps based on the previously reported protocol.^16^

### KrakenFISH for sequential, multi-condition smFISH

KrakenFISH is a modified version of previously reported AutoFISH, an open-source toolbox for automated, cost-effective sequential smFISH experiments. Briefly, the system consists of an automated fluidic system that is mounted onto a microscope and enables automatically alternating buffer exchanges with widefield fluorescence imaging rounds. The system was modified by addition of Luer connected tubing and a multi-way, computer-controlled valve to be compatible with 6 channel Ibidi microslides. Our hybridization, washing and imaging buffer recipes and the open-source building plans and software control packages are described in detail in a previous publication^16^ and GitHub (https://github.com/fish-quant/autofish/). Primary oligos were bioinformatically designed using our in-house software tool Oligostan.^27^ Primary oligo sequences consist of a portion complementary to a target RNA, and a barcode sequence specific to one RNA species of interest. The barcode sequence was added both at the 5’ and 3’ end of each oligo. Secondary oligos consist of readout regions that are complementary to the primary oligo barcode region of their corresponding gene and a sequence that is complementary to an imager probe. Our imager probes are conjugated to either Cy3 or Cy5 fluorophores, enabling imaging of two genes simultaneously each round. To remove readout probes from the previous rounds, we employ displacement probes complementary to a region larger than the barcode region of the readout probes. This modular probe design approach was designed to reduce recurrent experimental costs. The KrakenFISH setup reported in this project was mounted onto a Thunder Imager live cell (Leica microsystems, Germany) epifluorescent microscope. Image acquisition was performed in 5x5 tiles for each of the three conditions using 63X 1.4NA oil immersion objective (Leica microsystems, Germany), with 45 slices with a distance of 0,4μm between consecutive slices. Given cells that were not illuminated during live cell imaging were washed off, identifying the area in which live cells were imaged was straightforward, requiring locating clear patches of adhered cells.

### Live cell image processing and feature extraction

Live cell images consisted of 4x4 tiled brightfield images across 4 focus planes with a distance of 1,5μm between each slice, obtained for 21 timepoints spanning 7 minutes and 21 seconds. All analyses were conducted on a single focus plane. For each condition, last timepoint images were computationally stitched with tile blending and segmented using a finetuned Cellpose model, followed by manual quality control to correct for segmentation errors. This initial segmentation was used to provide seeds for back-propagation of SAM2^18^ across all timepoints for cell tracking. Since stitching can result in artefacts due to blending of cell images in overlap region, to obtain a more precise representation of cells, each cell was re-mapped back onto an individual tile and re-segmented using a more precise Cellpose segmentation model finetuned through manual annotation. Annotated images were made to include fine features of cells such as lamellipodia and small protrusions. Together, this approach allowed to obtain both x,y positions for each cell centroid at each timepoint, as well as precise contours for each cell at each timepoint. In total, we extracted 163 features for each cell, describing cell neighbourhood, morphology, trajectory and combinations thereof. All features are described in Supplementary Table 1. Morphological features included descriptors such as cell area, perimeter length and elliptical features derived from elliptical Fourier descriptors (EFDs). This last category of features, originally defined by Kuhl and Giardina^28^, precisely captures cell outline characteristics at different spatial frequencies, with low-order coefficients reflecting global shape and higher order coefficients encoding fine contour variations. They are derived to compute magnitude and elliptical area because they are rotation-invariant, in opposition to EFDs. All these descriptors were temporally aggregated through mean, minimum, maximum and standard deviation to capture both the central tendency and variability over time. All features were normalized based on mean and standard deviation, and missing values were replaced by median value.

### Estimation of smFISH spots for each cell

Fluorescent smFISH images consisted of 5x5 tiled acquisitions with 45 planes in two channels for each hybridization round, except for the first round that included DAPI imaging to help better locate cells for segmentation. To avoid spot detection errors in merged areas, each tile was segmented separately, and each cell was assigned to a single field of view. Fixed cell smFISH images were segmented using a customised Cellpose model followed by manual quality control. smFISH spots representing individual RNA molecules were localized in images using FISH-quant.^19^ This involved establishing threshold levels for each RNA species using a semi-manual approach, whereby the “elbow method”^19^ was employed to provide initial threshold candidates that were then finetuned based on visual inspection. We used the same threshold for a given gene across experimental samples. RNA counts per cell were estimated by assigning located spots to cell mask boundaries for each RNA species. We used the same threshold for each gene across samples. Bright autofluorescent debris and aggregates are commonplace, particularly in non-adherent cell samples where these contaminants cannot easily be removed from cultures through media exchanges. We therefore developed a simple quality control step whereby a mask of these fluorescent contaminants could be generated through manual thresholding for each condition and cell segmentations overlapping with this nonspecific fluorescence mask could be automatically excluded from analysis.

### Matched dataset generation

Dataset scale differences (40X magnification for live cell imaging and 63X for smFISH imaging) were first adjusted for based on the known pixel size ratio. To find the exact overlap between the two image sets, we developed a metric consisting of the sum of the distances of each segmented cell in the last frame of the live cell time-lapse to the nearest cell in the segmentation of the smFISH dataset at a given overlap position between the two tiled segmentations. This metric underpins a brute-force algorithm designed to find the optimal translational shift minimizing the metric, thereby achieving a precise spatial alignment of the two masks. Upon spatial correlation, each cell in one dataset is paired with the closest matching cell from the other dataset. The robustness of this alignment is quantified through two metrics: the distance between the paired cells (d_1_) and the distance to the second closest neighbouring cell in the corresponding dataset (d_2_). We use these metrics to filter the matched cells based on the desired trade-off between matching certainty and number of matched pairs. For all results presented, we applied d_1_ = 100 pixels and d_2_ = 50 pixels. Cells without 21 timepoints or partially cropped cells were excluded from the dataset.

### Model training

Cells coming from both 24-hour stimulation samples were pooled for all subsequent analyses and only cells with at least two detected transcripts of CD8 were included in the analysis. To simplify the training task and ensure model interpretability, we employed a random forest classifier on the 163 live cell features to predict each gene as high or low expressed independently. Two models were trained differing in how gene expression labels were obtained. For the first model, gene expression thresholds were obtained manually comparing with non-stimulated naïve cell baseline expression levels. For the second model, only the one hundred lowest and one hundred highest cells were used to assess the performance of the model on edge cases. Hyperparameters for each gene were optimized using grid search, and performance was evaluated using k-fold cross-validation (k = 5). Because many genes exhibited substantial class imbalance, balanced accuracy was used as the primary metric. In all analyses, 20% of cells were withheld from training and used for testing.

## DATA AVAILABILITY

Imaging data for both smFISH and live cell imaging are available from the corresponding authors upon reasonable request.

## CODE AVAILABILITY

All relevant analysis code is available at the following link: https://github.com/solenebernard/IMAGENE

## ACKNOWLEDGEMENTS

We gratefully acknowledge the UTechS Photonic BioImaging (Imagopole), C2RT, Institut Pasteur, supported by the French National Research Agency (France BioImaging, ANR-24-INBS-0005 FBI (BIOGEN); Investments for the Future), and acknowledge support from Institut Pasteur for the use of the Thunder Imager live microscope. We also recognize with gratitude Institut Pasteur’s Direction des Applications de la Recherche et des Relations Industrielles (DARRI) for their funding of M.I.G. and C.S. through the Emergence Grant, and specifically thank Sierra Parker for her substantial support of this project. We also gratefully acknowledge the EIT Health Deep Tech Venture Builder program for their funding of S.B. This work was also supported by the Japan Science and Technology Agency (JST) CREST (24031865), the Uehara Memorial Foundation, and the Ministry of Education, Culture, Sports, Science and Technology (MEXT) of Japan, Fund for the Promotion of Joint International Research (International Collaborative Research) (24KK0103). Finally, are deeply grateful for support and insights provided by many our colleagues, particularly Jacques Bourg, Thomas Defard, Elric Esposito, Nassim Mahtal, Luis Romero Martin, Audrey Salles, Lesly Raulin, Valérie Monceaux, Nicolas Bellini, Daniel Krentzel and Hao Zhu.

## ETHICS DECLARATIONS

Competing interests

The authors declare no competing interests.

## Notes

### Competing Interest Statement

The authors have declared no competing interest.

