## Supplementary Material for "IMAGENE: Single-cell association of live cell imaging and gene expression profiles of non-adherent cells through photoactivatable adhesives"

### SUPPLEMENTARY FIGURES

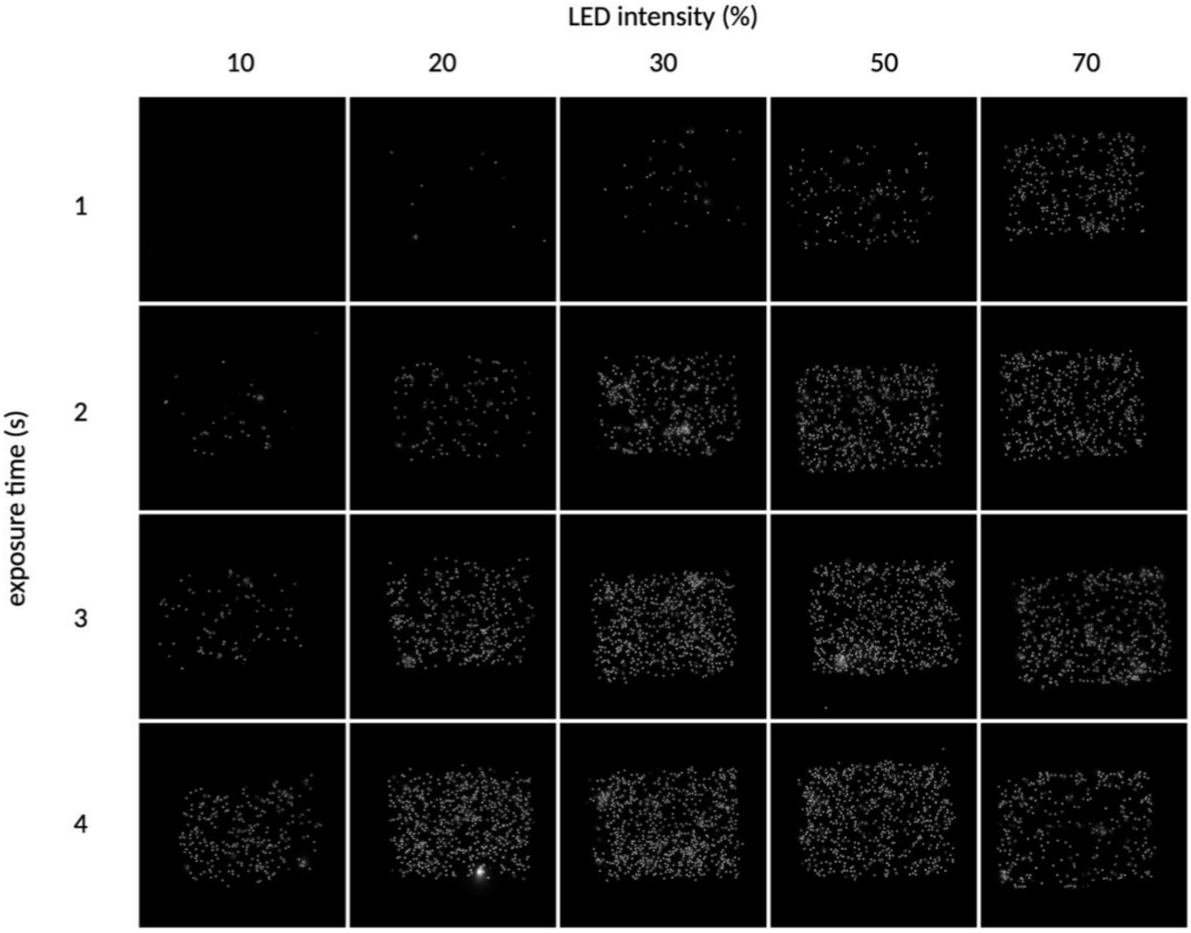


**Supplementary Figure 1. Photo-immobilization illumination parameter optimization.** Original illumination was done on one field of view using a 40x, 1.3NA objective and different illumination parameters. Resulting immobilised cell area was imaged in a 2x2 tiled image using the same 40x objective after washing, fixation and DAPI staining. Conditions of 50% illumination intensity and 2 seconds light exposure were selected for experiments.

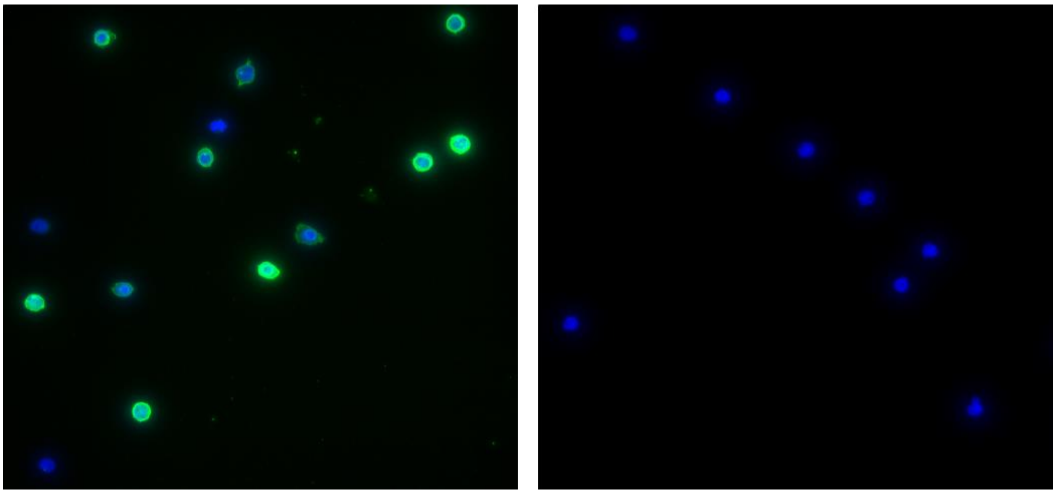

**Supplementary Figure 2. IMAGENE compatibility with immunofluorescence.** Bulk CD8+ T cells were isolated from frozen PBMCs, immobilized on PA-BAM using a 40x, 1.3NA aperture objective, fixed, blocked with bovine serum albumin (BSA) and stained with FITC-conjugated CD45RA antibody followed by DAPI (left). Control conditions were processed by incubation with blocking solution followed by DAPI (right). All samples were imaged using a 63x, 1.4NA oil immersion objective.

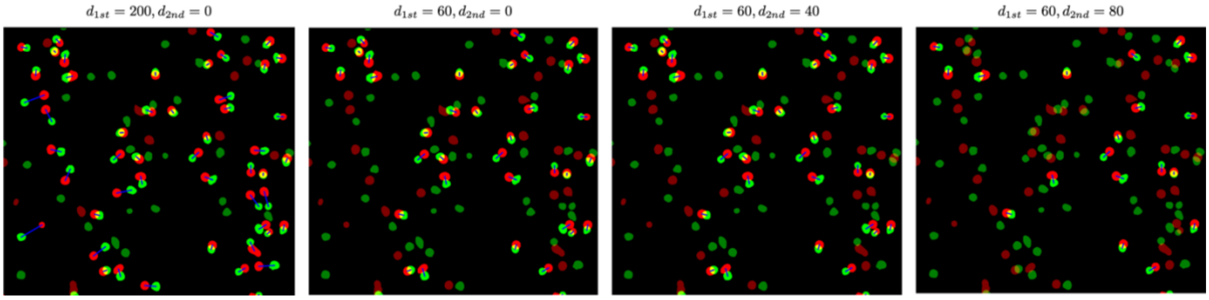


**Supplementary Figure 3. Influence of filtering parameters $d_1$ and $d_2$.** Visualization of the effect of tuning $d_1$ and $d_2$ (measured in pixels) on the resulting matches between segmented cells in brightfield (red) and segmented cells in smFISH (green). Matched pairs are displayed as bright and linked with a blue line, whereas unmatched cells appear as dim. The conditions are sorted from left to right with increasing restrictiveness.

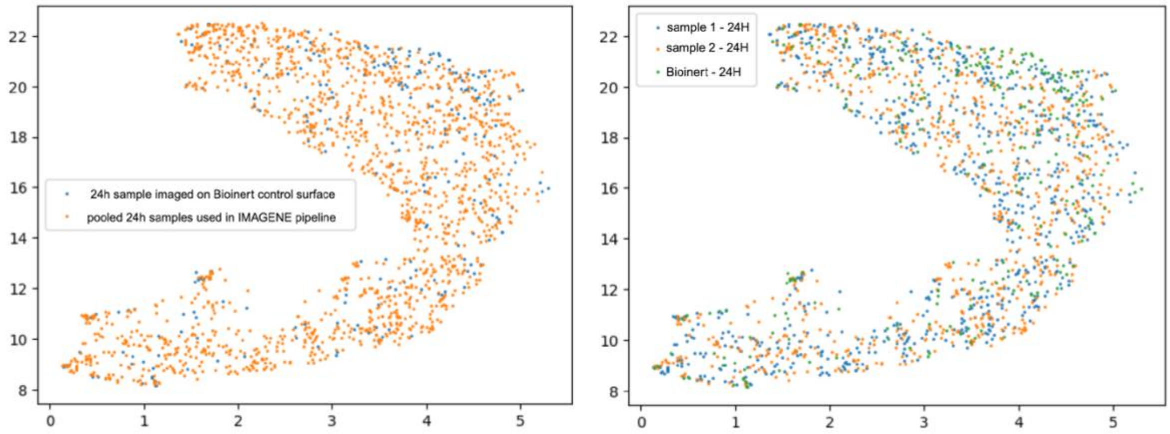


**Supplementary Figure 4. IMAGENE live cell imaging produces images comparable to common culture conditions.** (A) UMAP projection of the two 24 hour activated samples used in the IMAGENE pipeline (orange) versus a sample of cells from the same donor imaged on Ibidi Bioinert non-adherent microchannel slides (blue). (B) The same UMAP where all samples are color-coded separately demonstrating distributions of features are comparable between the two IMAGENE samples and between the IMAGENE samples and the samples imaged on the Bioinert substrate. Please note that while the same donor and stimulation condition was used for Bioinert and PA-BAM samples, cells in the Bioinert sample were used at a lower concentration.

**Supplementary Table 1.** Definition of all hand-crafted features. All areas and distances are computed in pixels. Features presented in bold represent time-dependent attributes that are aggregated using the mean, standard deviation, maximum, and minimum functions. This results in four distinct features for each time-dependent attribute.

| Feature name | Number of features | Description |
|---|---|---|
| **Elliptic descriptors from k=0 to 19.** | 20*4 = 80 | For a contour x(t) and y(t) being the periodic functions representing the x and y coordinates of a cell contour, the Fourier coefficients of order k named $a_k, b_k, c_k$ and $d_k$ are defined to solve the following equalities:<br>$x(t) = a0 + \sum_{k=0}^{+inf} x_k(t)$<br>$y(t) = c0 + \sum_{k=0}^{+inf} y_k(t)$<br>With<br>$x_k(t) = a_k cos(\pi kt) + b_k sin(\pi kt)$<br>$y_k(t) = c_k cos(\pi kt) + d_k sin(\pi kt)$<br>New descriptors, magnitude and elliptical area which are rotation invariant, are computed for each harmonic measuring respectively the size and the eccentricity of the ellipse k:<br>$\mathrm{Magnitude}_k = \sqrt{\lambda_{1,k}^2 + \lambda_{2,k}^2} \quad \mathrm{Ellipticalarea}_k = \lambda_{1,k}^2 \times \lambda_{2,k}^2$<br>Where $\lambda_{1,k}$ and $\lambda_{2,k}$ are the semi major and minor axes of ellipse of order $k$ obtained via singular value decomposition of the matrix, with $a_k, b_k, c_k, d_k$ being the Fourier coefficients previously introduced:<br>$M_k = \begin{pmatrix} a_k & b_k \\ c_k & d_k \end{pmatrix} = R_\theta \begin{pmatrix} \lambda_{1,k} & 0 \\ 0 & \lambda_{2,k} \end{pmatrix} R_\phi$ |
| **Absolute IoU** | 4 | The area of the intersection over union between the contour of the cell at timestamp t and t+1. |
| **IoU** | 4 | The area of the intersection over union between the contour of the cell at timestamp t and t+1, after centroid alignment. The feature only captures the change of morphology from t to t+1, without considering the change of position of the cell (as opposed to absolute IoU). |
| **Delta distance** | 4 | The distance between the centroid of a cell at timestamp t and t+1. |
| Total distance | 1 | The sum of all delta distance (distance between centroids at consecutive frames) across all timepoints. |
| Net displacement | 1 | The distance between the centroid of a cell at timestamp 0 and T=21. |
| Straightness | 1 | The ratio between net displacement and total distance. It falls between 0 and 1. |
| Mean turning angle | 1 | Average of absolute angles between vectors formed by the centroids of cell at t and t-1, with the vector formed by the centroids of the cell between t and t+1. |
| Convex hull area | 1 | The convex hull area of the path of the centroid of the cell with time, to capture the shape of its trajectory. |
| Area explored total | 1 | The total area explored by the cell during the time-lapse movie (obtained by overlapping all contours of cells). |

| | | |
|---|---|---|
| Alpha | 1 | Quantification of trajectory by the anomalous diffusion exponent $\alpha$.<br>The position $\mathbf{P}(t)P(t)$ of each cell at time t allows the calculation of mean squared displacement (MSD), defined as:<br>$\mathrm{MSD}(\tau) = \left\langle \lVert \mathbf{P}(t+\tau) - \mathbf{P}(t) \rVert^2 \right\rangle_t$<br>MSD typically follows a power law scaling:[29]<br>$\mathrm{MSD}(\tau) \propto \tau^\alpha$ ,<br>with alpha indicating deviations from classical Brownian diffusion with linear MSD growth. $\alpha$ =1 indicates Brownian diffusion, $\alpha<1$ sub-diffusive motion, and $\alpha>1$ super-diffusive behavior. |
| **Voronoi area** | 4 | Voronoi diagram "cell" (tile in the tessellated region) area for a given cell, calculated using all cell centroids. |
| **Distance boundary** | 4 | The closest distance from the cell to its closest boundary in the Voronoi diagram (which equals to half of the distance to its closest neighbour). |
| **N neighbours thresh k for k=100, 200 and 300** | 4*3 = 12 | The number of neighbouring cells located within a circle centered at the centroid of the cell and radius k. |

In addition to features described above, we also employed common features computed via scikit-image library[30] from the contour, like area, perimeter, eccentricity, equivalent diameter, bounding box area, extent, Feret diameter, major axis length, minor axis length, orientation and solidity, resulting in an additional 11x4=44 features. In total, we therefore obtained 163 features.
